# High-resolution bladder morphology in the intact mouse bladder: an intravital two-photon study

**DOI:** 10.1101/508879

**Authors:** Anna Schueth, Thomas L. Theelen, Sebastien Foulqier, Gommert A. van Koeveringe, Marc A. M. J. van Zandvoort

## Abstract

We developed a platform for intravital bladder two-photon microscopy (TPM) in healthy wild-type mice. It was our aim to make intravital bladder TPM simple and reproducible for researchers with access to TPM. In this study, the intact murine bladder was examined by means of autofluorescence (AF), second-harmonic generation (SHG), and different i.v. injected fluorescent dyes. All bladder layers with containing structures, such as cells, nerves, and vessels were detected in different colour-coded spectral channels, and shown in high-resolution images and real-time movies. The presented method opens up avenues for many future studies and applications, such as to reveal mechanisms and physiology in the natural state of the bladder. Researcher can use intravital bladder TPM to investigate events, such as migration and metabolism of (inflammatory) cells, nanoparticle uptake, or micro contractions in response to stimuli.

## Introduction

Worldwide, almost 600 million people are suffering from bladder dysfunctions, and therefore an increasing need for new treatments and diagnostic tools arises. Especially, in the field of imaging and microscopy many advances have been made and continuously new developments emerge. Bladder imaging is a very useful and important tool to investigate bladder pathology and the effect of treatments, both of the rodent bladder, as well as in humans (1, 2) (3, 4). Possible applications for imaging modalities are pathological changes in all layers of the bladder wall, which could be examined with a cellular or even subcellular resolution (5). Structural damages of the urothelium (6–9) or modifications of the muscle layer due to bladder outlet obstruction, and both detrusor under- and overactivity (10, 11) could be detected. Also the aged bladder shows ideal requirements for bladder imaging, and the increase in connective tissue and subsequent bladder fibrosis could be visualized, too (11).

Until the 20^th^ century, mostly classic imaging modalities, such as electron or bright field microscopy, which only provide 2D information, were applied to very thin, fixed bladder tissue sections.

To increase additional and new insights into the 3D bladder tissue architecture, it is important to choose an imaging modality that allows examination of thick slabs of fresh, living tissue in three dimensions. Intravital two-photon microscopy (TPM) is an excellent tool for real-time assessment of the physiological and metabolic state of the rodent bladder in its natural state. Sano and colleagues performed intravital FRET TPM on the bladder of transgenic mice, while inducing mechanical stretch to the tissue during the experiment (12). Since not all labs have access to expensive transgenic mice and special objectives and equipment, approaches for a simple, reproducible intravital bladder TPM protocol are needed. Here, we present an intravital bladder method with wild-type mice and a standard TPM water immersion objective. This allowed us to perform deep-tissue high-resolution imaging and we could visualize bladder morphology under nearly physiological conditions in real-time of the intact murine bladder.

Moreover, this intravital bladder TPM method may serve as an imaging platform for several applications to the rodent bladder in laboratories worldwide. Researcher can study immunology, pathophysiology and -morphology of the rodent bladder with detection of cell migration, influx of inflammatory cells and tracing of blood vessels using specific nanoparticles. Furthermore, micro-movement of tissue areas in different states can be studied at different depths in response to stimuli and inhibitors to elucidate signal transduction. Finally, technological advances of intravital bladder TPM bladder can lead to a micro-endoscopic two-photon imaging of human bladder for preclinical and clinical investigations.

## Materials and Methods

### Mouse model

Experimental protocols, approved by the animal ethics committee of Maastricht University, were carried out following institutional guidelines, and reported in accordance with the ARRIVE guidelines. All animals were obtained from Charles River (Wilmington, MA) and housed individually within a temperature-controlled environment with reversed 12-hour light/dark cycle, standard chow and water available ad libitum. In total, six adult C57BL/6 mice were used.

### Fluorescent probes

After warming, the mouse was positioned in a restrainer for i.v. injections. Subsequently the following markers were injected in a total volume of 200μl PBS: 1) SYTO 13 marker (2.5 μM, Life Technologies Europe BV, Bleiswijk, Netherlands) to stain cell nuclei and 2) yellow-green Fluoresbrite^®^ microparticles (size 1, 0 μm, 11,3 x 10^4^ beads/ μl, Polysciences Europe GmbH, Eppelheim, Germany) for blood vessel tracing. The markers were either used combined, or separately. The microparticles have an excitation maximum at 441 nm and an emission maximum at 486 nm, with Fluorescein being the used fluorophore. SYTO13 has an excitation maximum of 488 nm and an emission wavelength of 509 nm.

### Anaesthesia

Prior to surgery and after i.v. marker injection, the mouse was anaesthetised by injecting a combination of Ketamine (Ceva Tiergesundheit GmbH, Duesseldorf, Germany) (75mg/kg) and Medetomidine (Fort Dodge Veterinaer GmbH, Wuerselen, Germany) (0.5 - 1mg/kg) subcutaneously. Anesthesia was maintained during the entire procedure by additional injection every 30 min, while monitoring the vital signs and the depth of anaesthesia via the pedal reflex. The type of anaesthesia was selected based on minimal adverse effects on bladder function.

### Abdominal surgery

The mouse was kept (throughout surgery and the entire imaging experiment) on a heating plate (36-37°C) in a lateral position (Fig.1). After sterilizing (70% isopropanol) the shaved abdomen of the anaesthetized mouse, a 1-2 cm long medial abdominal incision was made, from posterior/caudal to anterior/rostral to expose the urinary bladder. The bladder was gently pulled out of the abdominal cavity, fixed with tissue adhesive (Histoacryl, B. Braun, Melsungen, Germany) to a metal stand and surrounded by agarose (Life Technologies Europe BV, Bleiswijk, Netherlands) and aquasonic ultrasound gel (Parker Laboratories Inc., Fairfield, NJ, USA) to keep the bladder moist and ensure the imaging process (Fig.1). At the end of each experiment the animal was sacrificed by cervical dislocation.

**Fig. 1.**
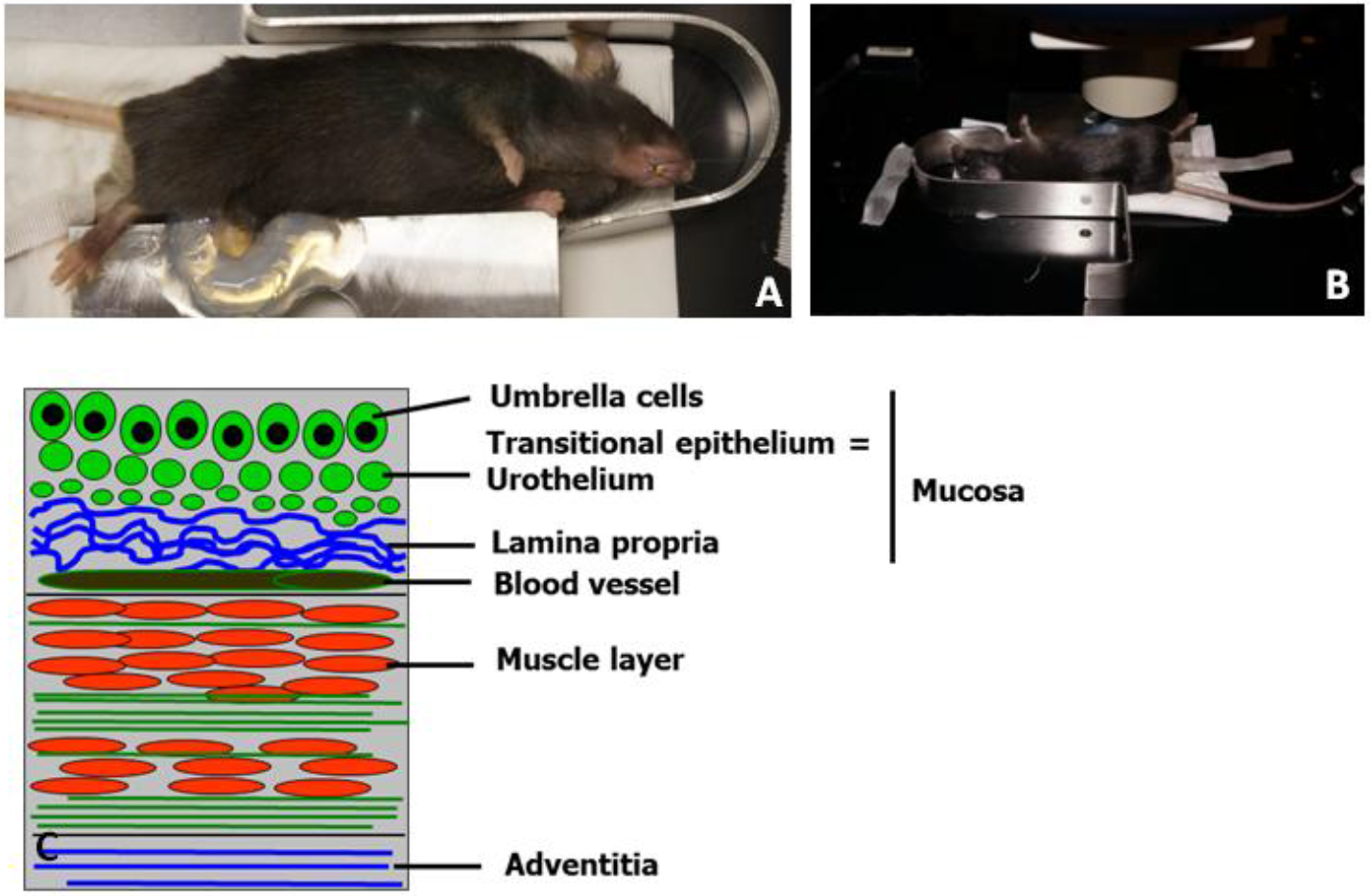
Experimental setup for intravital TPLSM bladder imaging with C57BL/6 mouse, after abdominal surgery and exposed urinary bladder. The anesthetized mouse is placed on a heated plate (36-37 degrees) and the bladder is stabilized with tissue adhesive and surrounded by agarose gel and ultrasound gel.

### In vivo TPM bladder imaging and imaging acquisition

TPM imaging (on Leica TCS SP5 MP, Leica Mikrosysteme Vertrieb GmbH, Wetzlar, Germany) was performed immediately after surgery. Ultrasound gel and physiological saline solution (0, 9 %, 154 mmol/l) (B.Braun, Melsungen, Germany) were applied on the adventitia/serosa of the exposed bladder to maintain humidity (Fig. 1).

Working distance of the objective was 2 mm, while the excitation source was a 140 fs-pulsed Ti:sapphire laser (Chameleon Ultra II, Coherent Inc., Santa Clara, CA, USA), mode-locked at 800-820 nm. To avoid photo-bleaching and tissue damage, laser power was kept between 30 and 50 % (100 %= 250-500 mW at the sample surface). Fluorescence emission was detected using photomultiplier tubes (Hamamatsu, R9624, Japan) in three wavelength ranges: 385-489 nm (blue, SHG of collagen), 489-563 nm (green, SYTO13, Fluoresbrite^®^ microparticles, AF of elastin, nerves), and 568-700 nm (red). Imaging settings were optimized for detecting both AF structures and stained structures. Image acquisition settings can be found in Table 1.

**Table 1.**
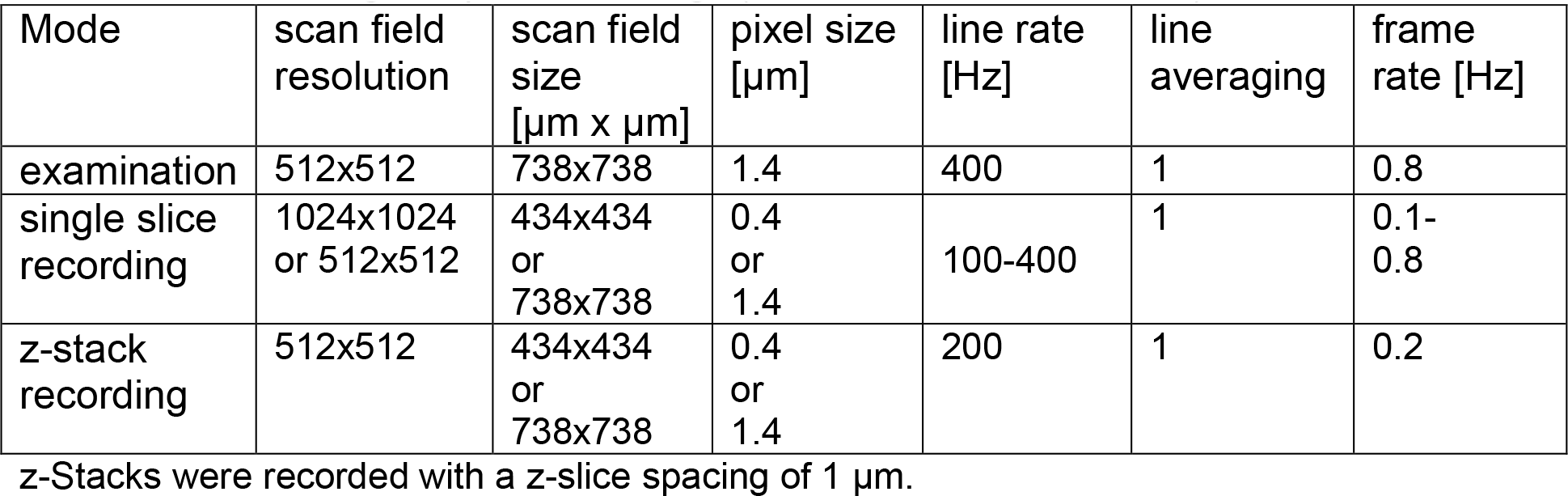
TPLSM image acquisition settings (without resonance scanner)

### Image data analysis and processing

Images were recorded and analysed with Leica Application Suite Advanced Fluorescence (Leica Microsystems). For further processing of the data, for instance 3D reconstructions both ImageJ/Fiji (13) (Rasband, W.S., ImageJ, U. S. National Institutes of Health, Bethesda, Maryland, USA, http://imagej.nih.gov/ij/, 1997-2012.) and IMARIS (Bitplane, Zurich, Switzerland) were used.

## Results

### Developing intravital bladder imaging in mice without resonance scanning

Although the bladder was stabilized (Fig.1), bladder movements as a result of contractions were still visible and indicating viability. The novel intravital TPM bladder set-up allowed imaging of the entire bladder wall in the living mouse for up to six hours.

### High-resolution bladder morphology in 2D and 3D in living mice

*In vivo* TPM bladder imaging of the complete and intact bladder of living wild-type mice, schematically given in Fig. 2A, revealed the typical layers and structures of the bladder wall (Fig. 2–4), by means of imaging of AF from intrinsic fluorophores, such as NAD(P)H, as well as i.v. injected markers. An orthogonal section through the complete bladder wall, imaged with intravital TPM, is given in Fig. 2B, indicating the various most prominent structures.

**Fig. 2.**
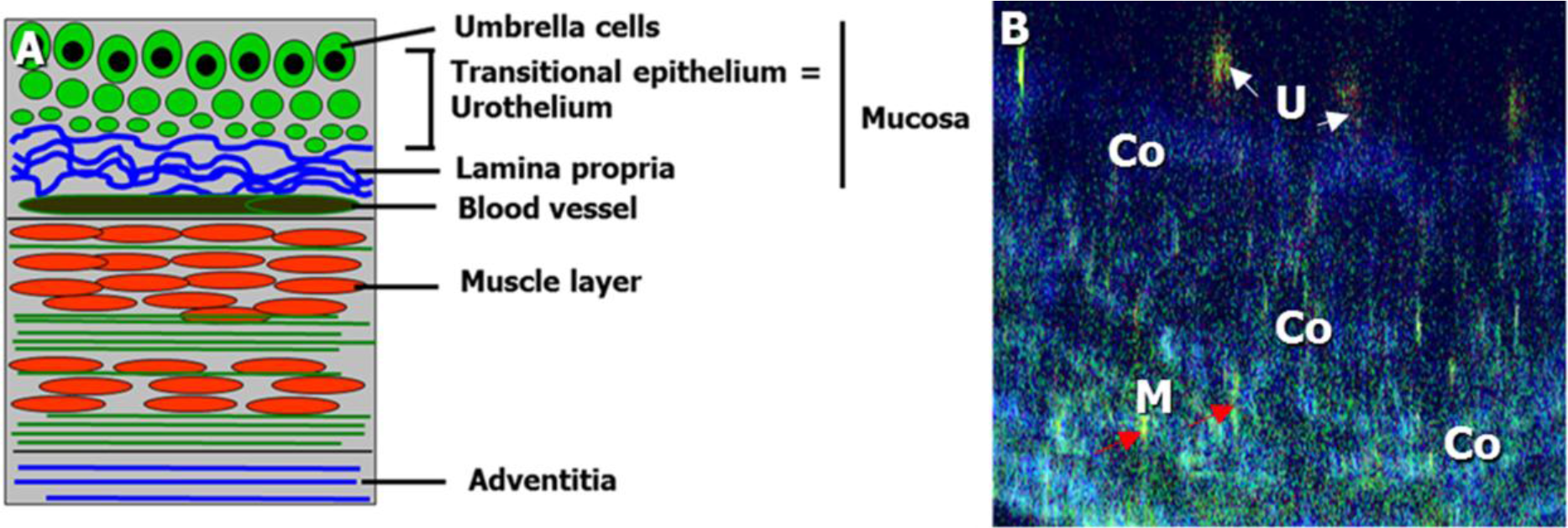
A) Schematic drawing of the urinary bladder wall, showing the different layers; the transitional cell epithelium (=urothelium), lamina propria (LP), muscle layer, and adventitia/serosa. The superficial layer of the urothelium is composed of the umbrella cells. Urothelium and adjacent LP together form the bladder mucosa. The muscular layer consists of several muscle bundles, built up by elastic fibers. B) Autofluorescent, label-free intravital TPLSM image, showing an orthogonal section of the complete bladder wall. The green urothelium is indicated by U and white arrows (NAD(P)H signal), the blue collagen layer (second-harmonic generation signal) is indicated by Co and the green muscle layer is indicated by M and the red arrows. Scale bar: 20 μm

**Fig. 3.**
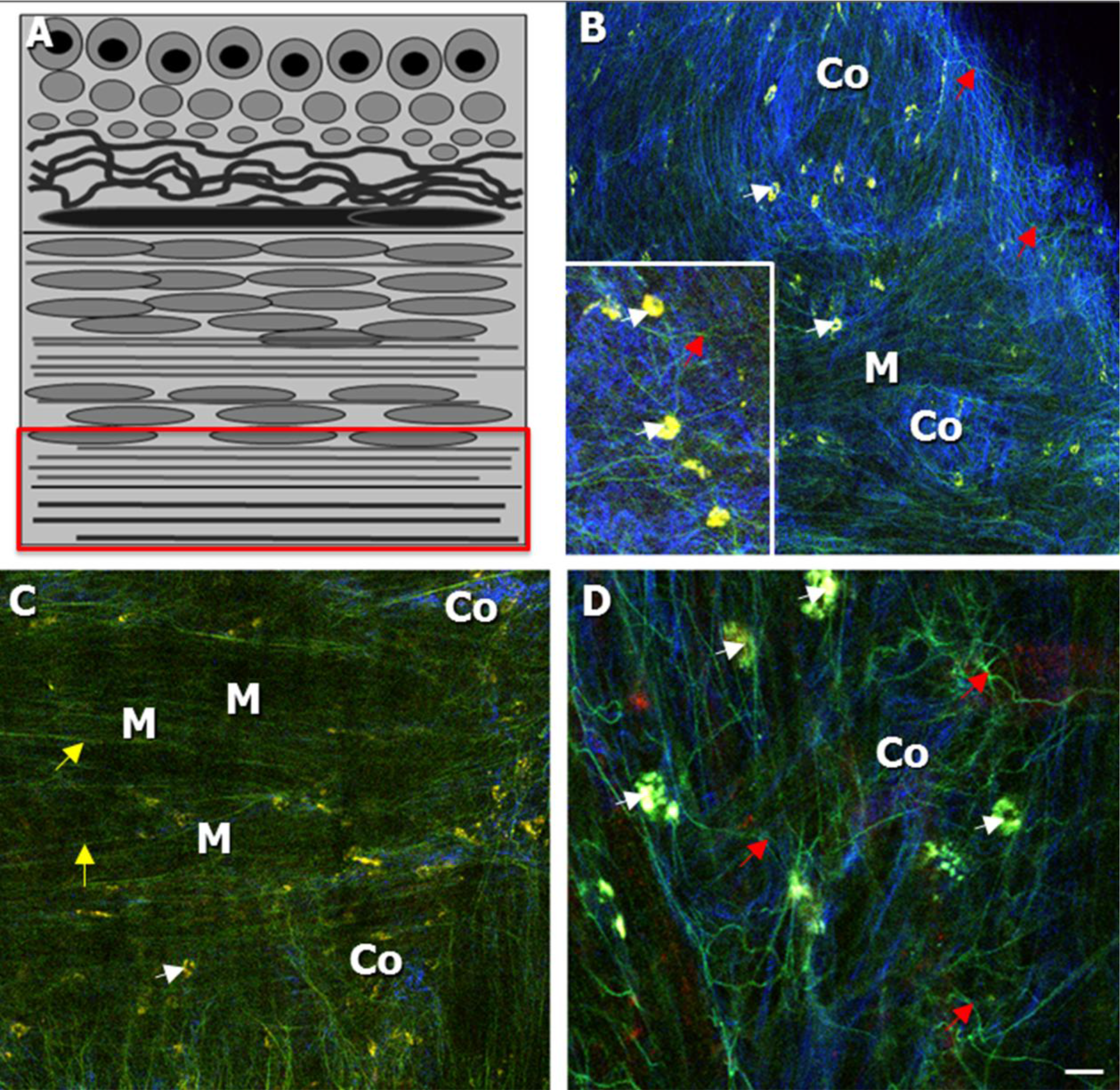
A) Schematic drawing of the bladder wall, indicating the imaging focal plane of B and C. B, C): Intravital TPLSM AF label-free images of the serosal collagen and muscle layer of the living bladder of healthy wild-type mice, while imaged via the adventitia. B) The serosal collagen layer (indicated by Co) was appearing in the second, blue, spectral channel (SHG signal), with nerves (red arrows) and macrophages appearing in the first, spectral channel (white arrows). Muscle bundles (indicated by M) were also partly detected, in the first, green spectral channel C) The muscle bundles of the muscle layer (indicated by M), were recognized due to elastic fibers (indicated by yellow arrows), running in both a longitudinal and circular direction and appeared in the first, green, spectral channel. D) For comparison, an *ex vivo* TPLSM bladder image, showing macrophages (indicated by white arrows) and nerves (indicated by red arrows) in the first, green spectral channel. Scale bar: 20 μm

**Fig. 4.**
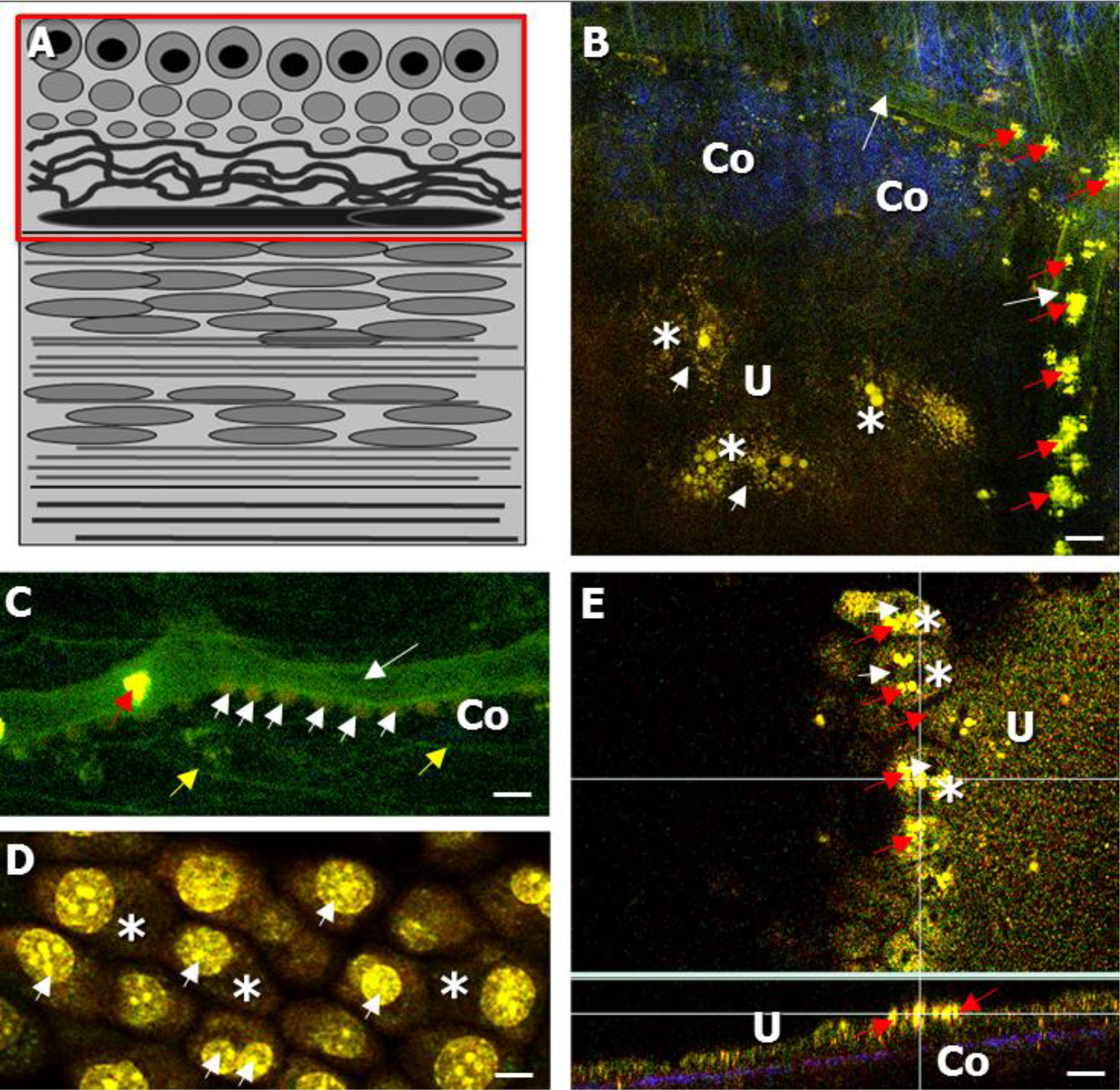
A) Schematic drawing of the bladder wall, indicating the imaging focal plane of B-D. BD): Intravital TPLSM AF images of the lamina propria (LP) and urothelial layer (U) of the living bladder of healthy wild-type mice, while imaged via the adventitia (i.v. injection of Fluoresbrite^®^ microparticles shown in B, C and SYTO 13 shown in C, D). B) The LP was containing collagen (SHG) and appearing in the second, blue spectral channel (indicated by Co). Blood vessels, with a diameter of 10-20 μm (white arrows) were seen in the first, green spectral channel and could be traced by means of microparticles (red arrows). The (sub) urothelium, indicated by U showed urothelial cells and umbrella cells with green fluorescent cytoplasm (white asterisks) and a dark-non fluorescent nucleus (white arrows). C) Blood vessels (white arrow) in the LP could be confirmed by SYTO13 staining of the nuclei of endothelial cells (white arrowheads). Also, elastic fibers were seen in the first, green, spectral channel (yellow arrows). D) Using SYTO13 staining the nuclei of the umbrella cells (white asterisks) could be detected due to a strong, green fluorescence (white arrowheads). E) The *ex vivo* imaging of the dissected bladder showed a distribution of the i.v. injected microparticles (red arrows) to the urothelial layer (indicated by U). The orthogonal section of the urothelium (Fig. 4 E, indicated by U) indcates an adhesive layer of microparticles (red arrows) and no particle uptake within the urothelial cells. Scale bar: 20 μm

While imaging the bladder adventitia/serosa (focal plane indicated in Fig. 3 A), a 2540 μm thick layer of collagen in the blue spectral channel (SC) (SHG, indicated by Co) was detected. The collagen layer contained AF macrophages, notable due to their typical shape and location (Fig. 3 B, insert, white arrows). Also, AF nerve fibers were observed in this layer, detected in the green SC (Fig. 3 B, red arrows). We note that the macrophages and afferent nerve fibers were both seen in our previous *ex vivo* study (Fig. 3 D, white and red arrows, respectively), where their origin was confirmed using immuno-histochemical-stainings (14). Already in this layer some muscle bundles were detected in green (Fig. 3 B, indicated by M).

Beneath the serosal connective tissue layer the muscle layer was found, (Fig. 3 C, indicated by M), which was characterized by the accompanying elastic fibers, in the green SC, to appear in a longitudinal or circular manner (Fig. 3 C, yellow arrow).

Below the muscle layer the lamina propria (LP)/ sub-urothelium (Fig. 4 B, C) could be visualized at a depth of approx. 25-50 μm. The imaging focal plane is indicated in Fig. 4 A. The LP contains collagen (Fig. 4 B, SHG, blue, Co), as well as elastic fibers and blood vessels (Fig. 4 B, yellow arrow, Fig. 4 B, C, white arrow, respectively, both green). The detected vessels ranged in diameter from 10 to 20 μm (Fig. 4 B, C, white arrow) and were confirmed with i.v. injected SYTO 13, which was staining the nuclei of the endothelial cells aligning the blood vessels (Fig. 4 C, white arrowhead). Fluoresbrite^®^ microparticles were taken up by the blood vessels and could be visualized in the green SC due to their yellow-green fluorophore fluorescein (Fig. 4 B, C, red arrow). The microparticles thus appeared very useful for tracing the blood vessels in the living bladder.

The transitional epithelium (=urothelium), which is located underneath the LP, (Fig. 4B, green, indicated by U, D) appeared due to the NAD(P)H signal of urothelial cells, especially in the superficial layer of the urothelium (Fig. 4 B, remove in situ D, white asterisk). While performing label-free AF intravital TPM bladder imaging, the nuclei of the umbrella cells (Fig. 4 B, white arrowhead) appeared dark with surrounding fluorescent cytoplasm. The thickness of the urothelial layer was observed to be up to 45 μm.

Additionally, our data allowed the production of z-stacks/movies, showing the entire bladder wall of the living mouse.

## Discussion

### Developing intravital bladder imaging in mice

Our novel intravital TPM bladder imaging setup enabled the visualization of the entire urinary bladder wall of living wild-type mice, under nearly physiological conditions. Depending on the natural urine filling state of the bladder, which is at most 0, 1 ml, we could perform imaging up to a depth of 200 μm. Stabilizing the bladder allowed for the acquisition of high-resolution images without additional triggering, while bladder contractions were indicating viability during the complete experiment. Moreover, no surgery was performed on the bladder itself and the anesthesia had no significant impact on bladder function. The experiment was less invasive than other existing intravital TPM experiments (15), because the bladder was kept intact. Also, the here presented bladder TPM protocol provides ideal conditions to examine the healthy bladder in vivo.

### Identification of label-free and stained bladder structure

TPM, being a color-coded imaging process revealed characteristic information in the three different colored SC: 1) AF, green 2) SHG, blue and 3) red. SHG enables the visualization of signals from the unique triple-helix structured collagen using excitation wavelengths of 800-820 nm (16).

In our previous *ex vivo* TPM bladder imaging study we could identify all layers of the murine bladder wall and structures therein: the adventitia/serosa, the muscle layer, the LP and the urothelium (= transitional cell epithelium) (14). In the current study, this typical morphology of the bladder wall could be detected *in vivo*, using both natural tissue AF, and i.v. injected fluorescent probes. High signals of auto-fluorophores, such as NAD(P)H, collagen and elastin, indicate a high cellular metabolism, and allowed the examination using SHG and AF imaging at excitation wavelengths between 800-820 nm (emission of NAD(P)H between approx. 380 and 510 nm) (17). In addition to intrinsic fluorophores, both the organ shape, as well as the contained type of collagen influences the AF imaging (18). While examining the filled bladder from the adventitia “outer side”, we detected a 25-40 μm thick layer, of connective tissue, in the blue SC. In comparison, serosal connective tissue of the dissected, emptied and stretched bladder has been shown to have a thickness between 15-25 μm (“ex *vivo* TPM bladder imaging”) (14). It has been proposed that the thickness of the collagen layer depends on the age of the experimental animal as well as the stretch status of the bladder. However, it is challenging to compare layer thickness between an *ex vivo* setup, where the bladder has been dissected and *in vivo* imaging in the living animal. The stretched status of an *ex vivo* bladder allows the visualization of all bladder wall layers, showing a clear separation between each layer. In contrast, using the complete, intact bladder in the *in vivo* imaging setup, with healthy bladder contractions and varying natural urine filling state does not allow the detection of the bladder wall layers to be completely separated from each other. In our opinion, this is more representative of the *in vivo* dynamic state of the bladder. Although the amount of collagen in the living mouse appeared to be increased compared to that in *ex vivo* isolated bladder tissue parts, it remains the same in an aged matched mouse (19).

Nerve fibers were found within the connective tissue, appearing in the green SC and therefore clearly distinguishable from the blue collagen. Moreover, nerve fibers could be differentiated from elastic fibers, appearing both in the green SC, using red Sulforhodamine B elastin staining (14). Nerves appeared in a rather curly shape, in contrast to the more straight-shaped elastic fibers, which build up the muscle bundles. Additionally, the collagen was spotted with macrophages, also appearing in green. Both macrophages and afferent nerves were confirmed in previous *ex vivo* work using immuno-histochemical-staining (14). The muscle layer, detected in green, appeared at a deeper level and was partially intermingled with the collagen layer and spotted with macrophages. Furthermore i.v. injection of yellow-green Fluoresbrite^®^ microparticles enabled blood vessel tracing in the muscle layer.

Finally, the LP and (sub) urothelium were detected in both the green and blue SC. The urothelium of the bladder wall showed a thickness of 45 +/- μm, while using our previous *ex vivo* imaging set up revealed a thickness of 30 +/- μm (14). This suggests that the urothelial thickness is highly dependent of the filling state of the bladder and its stretching status (20).

### Intravital TPM imaging in future urological research

Intravital TPM is expected to further evolve in the near future, in both basic as well as in clinical research. The here presented platform for intravital bladder TPM can be the basis for different approaches to study bladder function and physiology in mice. Recently, TPM has provided insight into inflammatory processes, such as lung fibrosis (21). Likewise, our technique may allow following bladder inflammation processes in time, even over the course of several weeks with the suitable animal model. Moreover, in the kidney microvascular function and drug delivery routes have been studied using TPM imaging (22). Recently, TPM intravital imaging has been performed in rat kidneys to reveal calcium dynamics in the proximal tubules (23). The application of *in vivo* calcium imaging to the bladder will enable the visualization of micro contractions within the bladder wall and provide information about the difference between local and spread effects of substances (through the entire bladder wall or specific regions).

In urology, *in vivo* imaging might be suitable for drug uptake studies, and the examination of both the permeability and drug response of layers, such as the urothelium and the detrusor muscle.

*In vivo* TPM is currently already used as a diagnostic tool in clinical Dermatology practice to study age-related changes of skin and its pathological state, for instance to detect melanomas or wounded skin (24). Furthermore, a combination of a multiphoton endoscope, together with the “DermaInspect” has been used to examine collagen and elastin of the (diseased) skin of patients, by only using SHG and AF(25). Adapting this technique to the bladder, could enable deep-tissue imaging of urothelial cell carcinoma and the detection of muscle infiltration without the need to resect the tissue. Moreover, this could be helpful in guiding a diagnostic biopsy and providing a surgeon with valuable tissue information intra-operatively in a shorter period of time.

It is crucial for further developing intravital TPM bladder imaging, to be applied via fiberoptics on human tissue in a pre-clinical set up, that no damage is caused to the examined tissue. Since we could not detect a drastic damaging effect on the tissue in our study, technological advances might enable the prototyping of a miniature bladder two-photon endoscope, like it has been developed for the kidney recently (26).

## Conclusions

With our *in vivo* TPM bladder imaging set-up we could visualize the complete and surgically intact bladder wall of the healthy, living wild-type mouse up to an imaging depth of 200 μm: from the outer “adventitia” to the inner “urothelial” side of the bladder wall. Stabilizing the bladder allowed for the production of high-resolution images, with healthy bladder contractions still indicating viability. By AF from intrinsic fluorophores, SHG and fluorescent probes we could detect several layers and structures, such as connective tissue, muscle bundles, and nerve fibers without labeling. Using microparticles, we could trace blood vessels. This is the first study describing intravital TPM bladder imaging, and shows that this technique bares the potential for a diagnostic use, such as a micro-scaled two-photon endoscope in urological patients in the future.

## Acknowledgements

The research was funded by the TRUST FP7-Marie Curie programme. The authors thank Jeroen Hameleers for technical support and Dr. Remco Megens for scientific advice.

